# Learning RNA secondary structure (only) from structure probing data

**DOI:** 10.1101/152629

**Authors:** Chuan-Sheng Foo, Cristina Pop

## 1 Introduction

The structure of an RNA molecule is critical to its function. Structured regions in an RNA molecule permit or impede the binding of proteins and small molecules, resulting in downstream effects on gene expression [24, 16]. However, accurate determination of RNA structure is an involved process, requiring tedious experimentation [8, 1, 10] or careful analysis of evolutionary constraints [9]. As such, several computational methods have been developed to determine RNA structure in silico. In this work, we focus on methods that predict an RNA’s secondary structure - the set of complementary Watson-Crick basepairs. These methods may be broadly classified into energy-based models [27, 14, 19], that compute a minimum free-energy secondary structure using experimentally-derived energies, and statistical models, which rely on data from a training set of sequences and their known structures in order to learn a model of secondary structure. In general, statistical methods for RNA secondary structure prediction outperform energy-based methods [20, 18]. However, their performance is found to be limited by the availability of training data [20].

The advent of high-throughput sequencing has led to the development of several genome-wide RNA structure-probing assays [26, 11, 23, 13, 5, 21] which could help plug this data gap. Such assays reveal which nucleotides are paired and which are not, but do not provide complete secondary structures. As such, existing statistical methods are unable to utilize these data for model learning. In this work, we present CONTRAfold-SE, which extends the statistical model of CONTRAfold [6] to model the structure-probing data as observations of possibly unknown secondary structures. This model can then be learned from datasets containing only structure-probing data, or a mix of known structures and probing data. We train CONTRAfold-SE on various combinations of structure probing data and complete structures and find that while genome-wide structure probing data provides modest improvement in prediction performance, with sufficiently dense probing data alone it is possible to learn a model that approaches the performance of energy-based methods. CONTRAfold-SE may be obtained from https://github.com/csfoo/contrafold-se.

## 2 Methods

### Modelling structure-probing data as observations of secondary structure

Our model assumes that structure-probing data are available at per-base resolution – that there is a probing signal for some set of bases in a given RNA sequence. While our probabilistic framework allows the use of any distribution in the exponential family to model the (processed) probing signal at each base, we chose the Gamma distribution since it is flexible in modelling the positive, continuous, and unbounded probing signal. In addition, the Gamma distribution fits the data well, and has been previously used to model signal from various probing assays [2]. We also assume that bases are independently modified, so that the resultant probing signals are independent of the actual location within the RNA sequence. However, as the reactivities of different bases to the probing mechanism could differ based on their identity and whether they are paired (for instance, DMS preferentially modifies unpaired adenines and cytosines), we have incorporated a separate distribution for each combination of base identity (A, C, T or G) and pairedness state (paired or unpaired) for a total of 8 separate Gamma distributions in our data model. Formally, for an RNA sequence *x* of length *L* with secondary structure *y* and associated structure-probing data *d* = (*d*_1_, …, *d_L_*), the distribution for the probing signal *d_k_* at base *k* in the sequence is given by

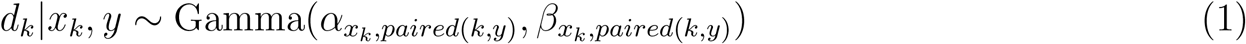

where *x_k_ ∈* {*A, C, T, G*} is the identity of base *k* in the sequence, *paired*(*k, y*) denotes whether base *k* in structure *y* is paired, and the Gamma density is defined as 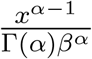 exp(*≤x/β*),for *x ∼* Gamma(*α, β*).

### The CONTRAfold-SE model and parameter estimation

#### Model specification

Let *x* be an RNA sequence of length *L* with structure *y* and *S* associated structure-probing datasets *d*. We denote by 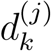 the probing signal for the *j*th data source at base *k* in the sequence. CONTRAfold-SE models the conditional joint probability of structure and probing data given sequence as

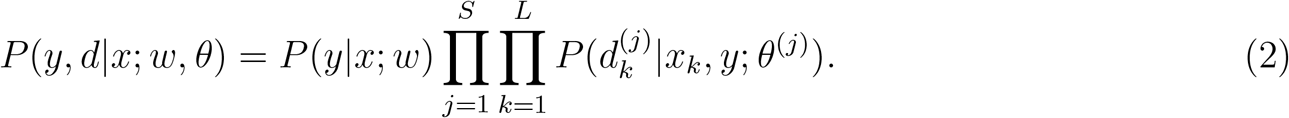

Here, the structure model *P* (*y|x*; *w*) is given by the conditional log-linear model of CONTRAfold with parameters *w*, and 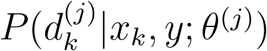 is the Gamma distribution as defined in Equation 1, with *θ*^(*j*)^ being the vector of parameters for the 8 Gamma distributions for dataset *j*. In the absence of structure-probing data, the CONTRAfold-SE model reduces to the CONTRAfold model.

#### Parameter estimation

Given a training set, we estimate the parameters, *w* and *θ*, by maximizing the conditional log-likelihood. Formally, for a training set *𝒟* = *𝒟_S_ ∪ 𝒟_𝒫_ ∪ 𝒟𝒮*+*𝒫* of sequences with: i) only known structures and no probing data (*𝒟_𝒮_*), ii) only probing data but unknown (missing) structure (*𝒟_𝒫_*), and iii) both known structure and probing data (*𝒟_*𝒮*+*𝒫*_*), we find *w*, *θ* that maximize the (regularized) conditional log-likelihood

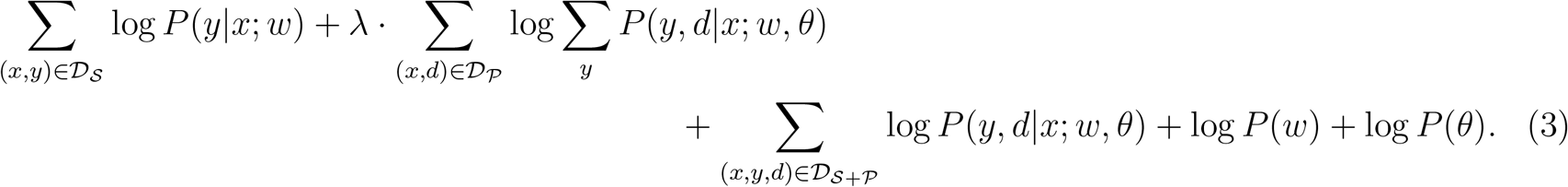

The hyperparameter *λ* (set by cross-validation) controls the weighting of data-only training instances as compared to instances with known structure, thus mitigating the adverse effects of noisy, partial data on model estimation; this strategy is common in the machine learning literature [17]. We used the L-BFGS algorithm [12] to find a local maximum of the likelihood. The key technical challenge is that the gradient computations for the second term in the sum (the likelihood for sequences with unknown structures) requires inference. Fortunately, the log-linear form of the structure model allows the data model to be represented as additional base-level features in the structure model. [7] independently presents a similar observation in the context of energy-based models for RNA structure, which have a similar log-linear form. We thus adapted the existing inference algorithms in CONTRAfold to efficiently compute the required gradients. While the log-likelihood for the CONTRAfold-SE model is non-convex, in practice our gradient-based parameter estimation algorithm achieves stable parameter estimates. A detailed description of the estimation procedure is found in Supplementary Methods.

## 3 Experiments

We evaluate CONTRAfold-SE in two scenarios: 1) with training sets of known structures augmented with data from genome-wide structure probing experiments, and 2) with a training set consisting solely of probing data. In each scenario, we determine the prediction performance of CONTRAfold-SE on TestSetA and TestSetB obtained from [20], which contain structurally disjoint sets of RNA structures from the RFAM database. We constructed training and test sets to minimize structural overlap in order to more accurately assess generalization performance. Details on the construction of these training datasets is included in the Supplementary Methods.

We evaluate methods based on the following per-sequence metrics averaged over the test set: sensitivity, positive predictive value (PPV; also known as precision), F-measure (harmonic mean of sensitivity and PPV), and accuracy. As CONTRAfold-SE, like CONTRAfold, offers a sensitivity-PPV tradeoff via a hyperparameter *γ*, which predicts more basepairs with increasing *γ*, we also compare the area under the sensitivity-PPV curve (AUC); for metrics other than AUC, we show the maximum value across varying *γ*.

### Dataset of known structures augmented with genome-wide probing-data

The training sets used in this scenario have two components making up 238 sequences: half with only known secondary structure and half with only structure-probing data. Probing-data only sequences are chosen to be the most data dense sequences across the probing datasets we use – the parallel analysis of RNA structure (PARS) [11] and DMS-seq assays [21]. We chose these genome-wide datasets as they were performed on the same set of (yeast) RNAs, allowing us to showcase the ability of CONTRAfold-SE to integrate multiple datasets, and explore the effects of such integration.

In the PARS assay, the RNA structure signal is obtained by treating RNA with enzymes that preferentially cleave either paired or unpaired nucleotides. The DMS-seq assay relies instead on the reactivity of unpaired nucleotides (primarily A and C) to the dimethyl-sulfate chemical; the DMS-seq assay was applied to both renatured RNA and live yeast. We denote these sources PARS, DMS-vitro, and DMS-vivo, respectively. We compare CONTRAfold-SE to CONTRAfold trained only on the set of sequences with known structures.

We find that while including a single source of structure-probing data has minimial effect on prediction performance compared to training on known structures alone, we obtain an improvement when using the two in-vitro data sources (Table 1).

**Table 1:**
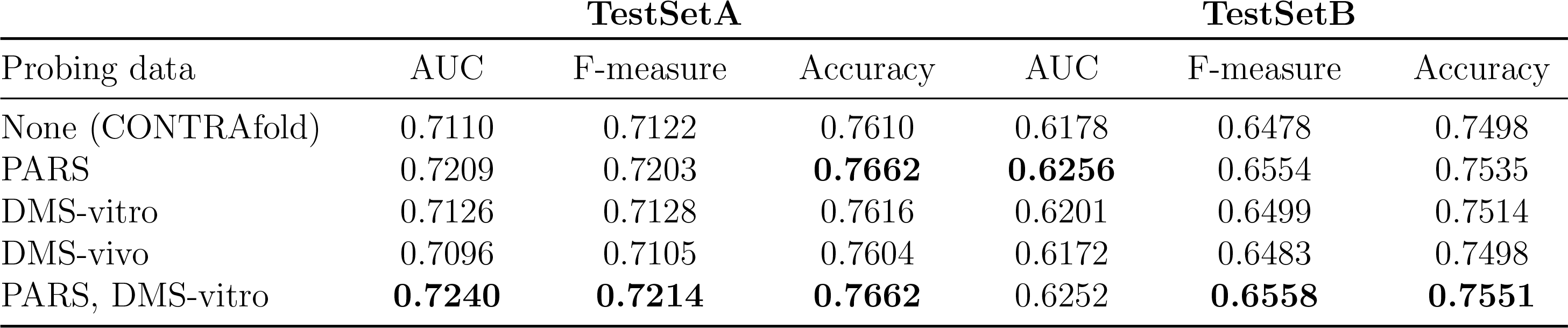
Performance on probing-data augmented training dataset.

### Dataset of only probing-data from RMDB

We next explored the possibility of learning a structure model using structure-probing data alone. As we and others [25] have observed that the PARS and DMS-seq genome-wide datasets we used in the previous section are sparse and have low signal-to-noise, we used datasets obtained with the method of selective 2-hydroxyl acylation analyzed by primer extension (SHAPE) [15] for structure-probing instead. This data is obtained from a single RNA species at a time, resulting in higher data density, and is also thought to be less biased [7]. We collected 17000 training sequences probed with the SHAPE method from the RNA Mapping DataBase (RMDB) [3] that had a high signal-to-noise ratio, and were not structurally similar to the test sets. We tuned the regularization hyperparameter (for L2-regularization on *w*) and performed early stopping using as a validation set the other test set not used for evaluation (*i.e.* TestSetA was the validation set when evaluating on TestSetB and vice versa).

While CONTRAfold-SE performs worse than UNAfold [14] (Table 2), a widely-used energy-based prediction method, this result was achieved *without the use of known structures* in the training set. In addition, comparing the learned parameters, we see that they are highly similar to those from a CONTRAfold model trained with known structures (Figure 1).

**Table 2:** Performance when model is learned only with structure probing data. We report sensitivity (Sens), positive predictive value (PPV) and the F-measure (F1).

| TestSetA | Sens | PPV | F1 |
| --- | --- | --- | --- |
| CONTRAfold-SE | 0.47 | 0.44 | 0.46 |
| UNAFold | 0.56 | 0.49 | 0.52 |

| TestSetB | Sens | PPV | F1 |
| --- | --- | --- | --- |
| CONTRAfold-SE | 0.53 | 0.45 | 0.49 |
| UNAFold | 0.61 | 0.48 | 0.53 |

**Figure 1:**
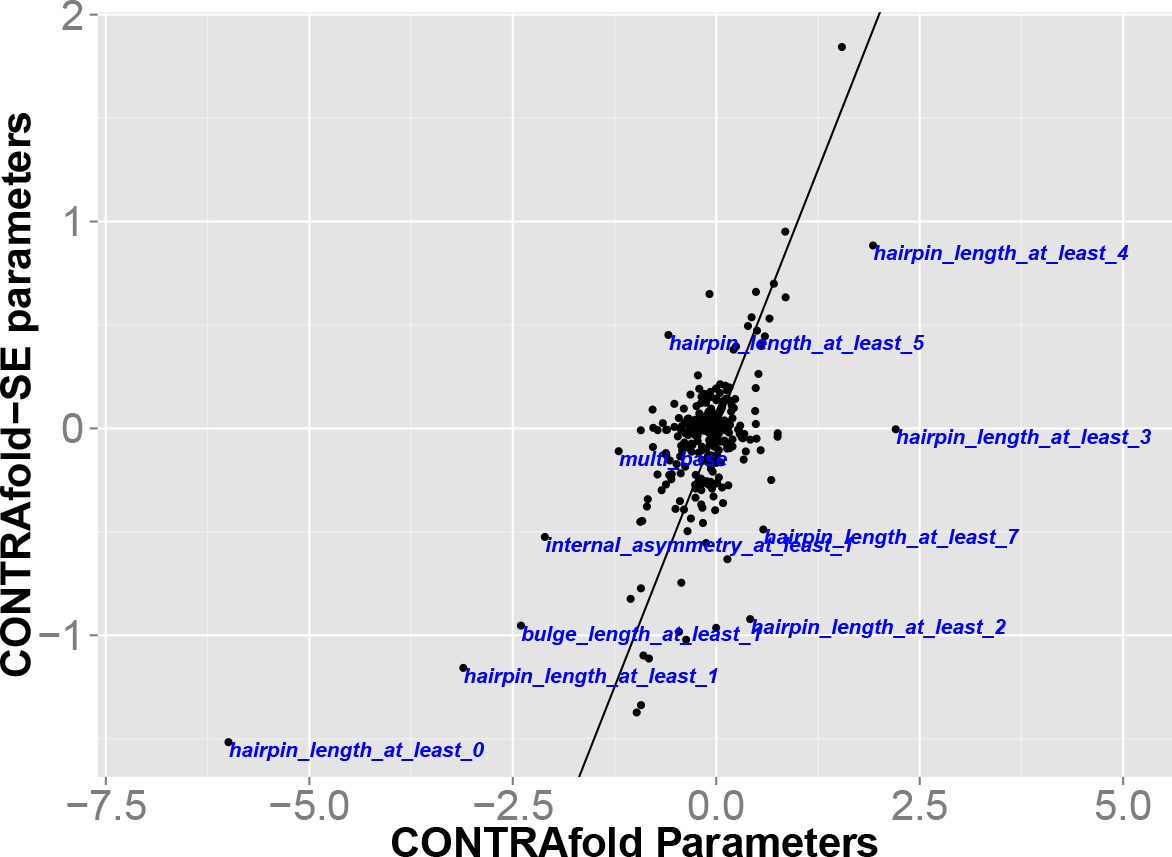
Comparison of parameters learned on structure-probing data alone (CONTRAfold-SE) with parameters learned on known structures (CON-TRAfold).

## 4 Discussion

We showed how structure-probing data can be used to learn models of RNA secondary structure. Our principled probabilistic framework enables easy integration of multiple probing datasets and handles situations where probing data or known structures are not available. While a similar approach has been used for improving secondary structure predictions with multiple probing datasets [22], the focus of that work was to model the characteristics of the probing data, and not to improve the underlying structure model. That CONTRAfold-SE is able to approach the performance of UNAfold when trained only on sequences with structure-probing data (without known structures) suggests that large collections of structure probing data could potentially be used to complement existing structures for training more sophisticated models. Finally, the sequences in the RMDB dataset we used were short (at most 103bp), and do not comprise a diverse set of RNAs. We speculate that with high-quality probing data from a diverse set of RNAs of various lengths, CONTRAfold-SE will be able to surpass the performance of energy-based models, and possibly that of statistical models. Incorporating the dependence of the SHAPE probing data on local structure context [4] may also help close the performance gap.

## Acknowledgements

We thank Rhiju Das, Daphne Koller, members of the Koller lab, and Tom Do for useful discussions.

